# *C. elegans* males optimize mate-choice decisions via sex-specific responses to multimodal sensory cues

**DOI:** 10.1101/2023.04.08.536021

**Authors:** Jintao Luo, Arantza Barrios, Douglas S. Portman

## Abstract

For sexually reproducing animals, selecting optimal mates is essential for maximizing reproductive fitness. Because the nematode *C. elegans* reproduces mostly by self-fertilization, little is known about its mate-choice behaviors. While several sensory cues have been implicated in males’ ability to recognize hermaphrodites, achieving an integrated understanding of the ways males use these cues to assess relevant characteristics of potential mates has proven challenging. Here, we use a choice-based social-interaction assay to explore the ability of *C. elegans* males to make and optimize mate choices. We find that males use a combination of volatile sex pheromones (VSPs), ascaroside pheromones, surface-bound chemical cues, and other signals to robustly assess a variety of features of potential mates. Specific aspects of mate choice are communicated by distinct signals: the presence of a sperm-depleted, receptive hermaphrodite is likely signaled by VSPs, while developmental stage and sex are redundantly specified by ascaroside pheromones and surface-associated cues. Ascarosides also signal nutritional information, allowing males to choose well-fed over starved mates, while both ascarosides and surface-associated cues cause males to prefer virgin over previously mated hermaphrodites. The male-specificity of these behavioral responses is determined by both male-specific neurons and the male state of sex-shared circuits, and we reveal an unexpected role for the sex-shared ASH sensory neurons in male attraction to endogenously produced hermaphrodite ascarosides. Together, our findings lead to an integrated view of the signaling and behavioral mechanisms by which males use diverse sensory cues to assess multiple features of potential mates and optimize mate choice.

## INTRODUCTION

Mate choice behavior has a central role in the fitness and evolution of sexually reproducing species.^1, 2^ As such, it provides an ideal context for understanding how genes shape behavioral programs. It is also an important entry point for understanding how and why biological sex influences the structure and function of the nervous system, as mate-choice behaviors typically differ by sex. Historically, mate choice has been thought to be more important for females, whose “choosy” behavior balances more indiscriminate mating patterns of males.^1, 3^ However, recent work in several species has shown that these dynamics can be more complex, with both sexes making active decisions about which mates to choose.^4–13^

In the nematode *C. elegans*, a powerful model for understanding the neural and genetic control of behavior, little is known about mate choice behavior. In this species, two features critical for mating system dynamics differ from those of most animals.

First, XX individuals are not females but rather simultaneous hermaphrodites, as they produce a limited pool of self-sperm (roughly 300)^14^ during larval development. While these self-sperm are present, hermaphrodites can reproduce by self-fertilization or by cross-fertilization, if inseminated by a male. After their self-sperm are depleted, XX animals become functionally female^15^ and produce further progeny only when cross- fertilized. Second, XO (male) individuals are rare in natural populations, as self- fertilization produces >99.8% XX progeny^16^ and is the predominant mode of reproduction in the wild.^17^ For these reasons, mate-choice behavior is less relevant for young adult hermaphrodites^18^ (indeed, self-fertile hermaphrodites typically resist mating attempts, presumably because they favor selfing^19^). In contrast, mate choice might be particularly important for males, as they sometimes face few competitors but many potential mates.

Several signals have been implicated in the attraction of *C. elegans* males to hermaphrodites.^20^ The best characterized of these signals are the ascaroside pheromones, aqueous, non-volatile derivatives of the dideoxy sugar ascarylose. As their biosynthesis differs by sex, stage, and an animal’s physiological status, these molecules are thought to enable a modular system of communication for nematodes.^21–25^ Several ascarosides, particularly ascr#3, are produced more abundantly by hermaphrodites and serve to attract or retain males.^26^ Other studies have identified non-ascaroside sex pheromones that are produced by hermaphrodites and elicit male-specific responses.^27–,30^ While the molecular nature of these remains unknown, one class, the volatile sex pheromones (VSPs) is regulated by hermaphrodite reproductive status and has been proposed to allow sperm-depleted hermaphrodites to attract males.^27, 30^ Even less well understood is the potential importance of cuticle-associated cues, well-known in other invertebrates^31^ but little-studied in *C. elegans*. Previous work has indicated that glycosylation of cuticle-associated proteins on the surface of adult hermaphrodites is important for mate-recognition by males,^32^ but the nature of this signal remains unknown. Moreover, how males use these diverse signals to influence mate choice is unclear. Often, male responses to these cues have been studied in artificial contexts, for example, using hermaphrodite-conditioned media or chemically synthesized pheromones.^27–29, 33–35^ Thus, the contribution of these signals in their native context remains poorly understood (Figure 1A).

**Figure 1.**
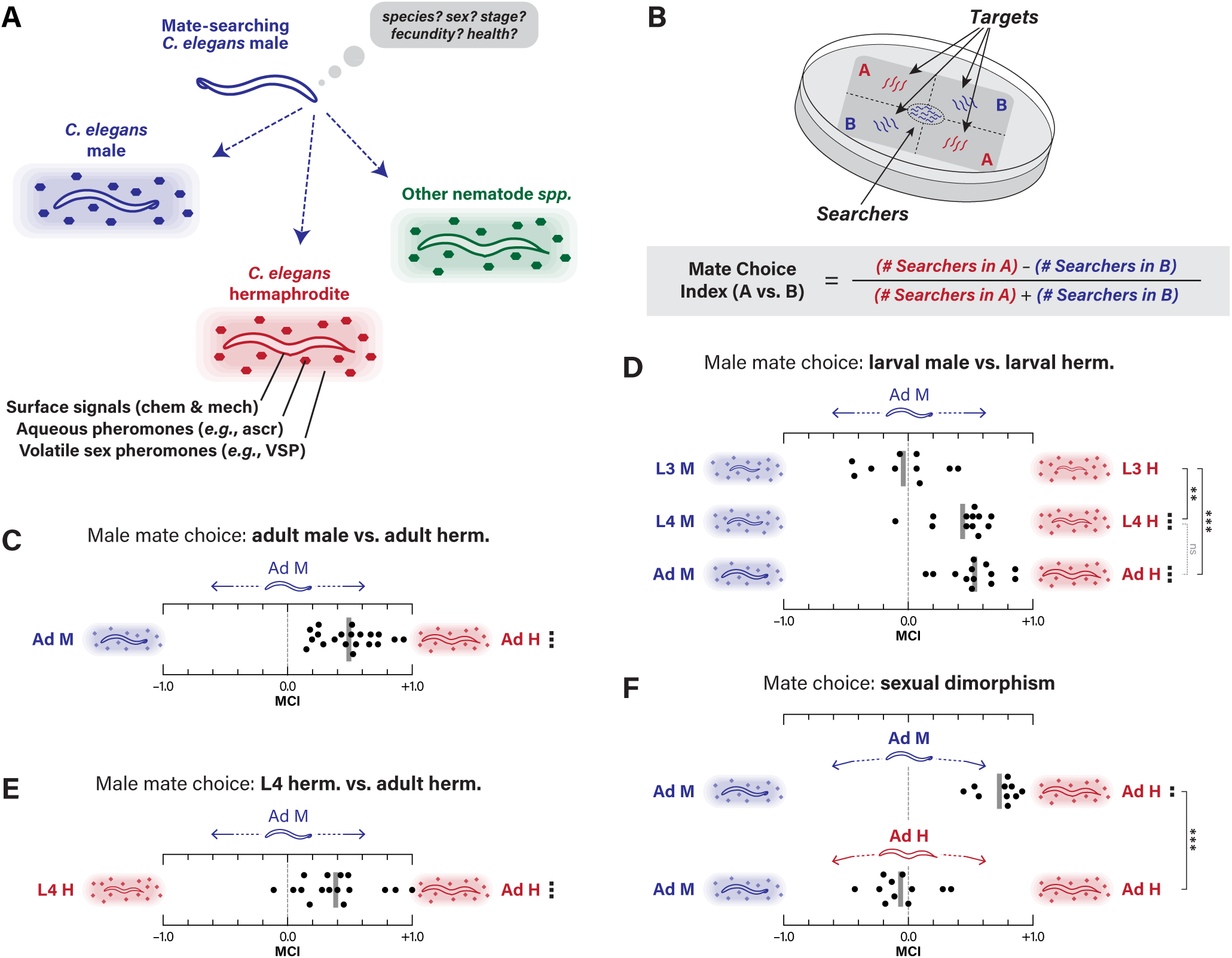
Mate-choice behavior is male-specific and allows males to distinguish the sex and developmental stage of potential mates. **(A)** In their natural environment, *C. elegans* males encounter adult conspecifics of both sexes, larvae, and other nematodes. Multiple classes of chemical and physical signals have been implicated in *C. elegans* mate recognition, including surface signals (indicated by the outline of the body), ascaroside pheromones (small colored hexagons), and volatile sex pheromones (VSPs, indicated by gradient shading around the animal). **(B)** The Mate Choice Assay (MCA). Immobilized targets of two classes are placed in two opposing quadrants; after 30 mins, searchers are placed in the center. Their positions over the next 90 mins are used to calculate a Mate Choice Index (MCI) (see Materials and Methods). **(C-E)** MCAs with control adult male searchers and (C) adult male vs. adult hermaphrodite targets, (D) male vs. hermaphrodite targets at L3, L4, and adult stages, and (E) L4 vs. adult hermaphrodite targets. **(F)** MCA with control adult male and hermaphrodite searchers and adult male vs. hermaphrodite targets. In this and all following figures, schematic diagrams show the nature of the cues potentially produced by each target class, with red indicating hermaphrodite and blue indicating male. Each data point represents an individual MCI derived from a MCAs using ∼10 searchers; the vertical gray bar indicates the mean MCI. For each tester-target condition, we carried out a Wilcoxon signed-rank test to ask whether the MCI was consistent with the null hypothesis that males had no preference for either targets class (*e.g.,* MCI = 0). Black squares to the right of each row indicate the *p* values associated with these tests: ▪, *p* ≤ 0.05; ▪▪, *p* ≤ 0.005; ▪▪▪, *p* ≤ 0.001. To compare MCI values between different classes of searchers, we used a Mann- Whitney test (to compare two genotypes) or a Kruskal-Wallis test with Dunn’s correction (to compare more than two genotypes). Asterisks indicate *p* values associated with these tests: \**p* ≤ 0.05; \*\**p* ≤ 0.005; \*\*\**p* ≤ 0.001. We only carried out tests necessary to evaluate specific pre-established hypotheses. Comparisons made whose outcomes were not statistically significant are shown in gray dotted lines and are labeled “n.s.”.

The strong motivation of adult *C. elegans* males to find mates is vividly illustrated by their tendency to abandon a food source in the absence of a suitable mate.^36^ This food-leaving behavior is almost completely suppressed in the presence of an adult hermaphrodite, but only weakly so by another adult male.^36^ Food-leaving resumes once conspecifics are removed, suggesting that a contact-dependent signal or a labile chemical cue is the primary cue for retention.^36^ Further studies highlighted a role for detection of contact cues by sensory neurons of the tail rays, male-specific sensilla that contact the hermaphrodite body during the first step of mating behavior.^37, 38^ These studies highlight the importance of physical contact in retention but leave open questions about the role of secreted signals and mechanisms that males use to evaluate the relative quality of potential mates.

Here, we use a simple choice-based social interaction assay to probe the features of potential mates that males assess, the signals that carry information about these features, and the mechanisms by which males detect and integrate them. Because behavior in this assay is evoked by endogenously produced cues presented at physiologically relevant levels in their native context, it provides a more ethologically meaningful setting in which to study male mate-choice behavior. We find that males show a strong preference for interacting with adult hermaphrodites over adult males, but hermaphrodites exhibit no such preference. By manipulating animals’ ability to produce and detect different categories of potential cues, we demonstrate that males use multiple, partially redundant features to distinguish the sex of potential mates.

Further, we find that males can distinguish the developmental stage of potential mates as well as their mating history (mated vs. virgin) and nutritional status (well-fed vs. starved). We find that ascarosides, VSPs, and surface glycosylation cues have distinct roles in these decisions, with ascarosides being important for the assessment of developmental stage, mating status, and nutritional status. Finally, we show that males’ ability to detect and integrate these cues depends on the male state of sex-shared neural circuits, such that masculinization of the hermaphrodite nervous system is sufficient to elicit male-like mate choice behavior. At least two sex-shared neurons, ADF and ASH, have central roles in male mate choice. Together, these findings reveal a multimodal logic by which males use diverse sensory cues to assess multiple features of potential mates and make optimal mate-choice decisions.

## RESULTS

### Adult *C. elegans* males robustly distinguish the sex of potential mates

In animals without courtship rituals, mate choice behavior can be difficult to disentangle from other aspects of reproduction. To study mate choice behavior in *C. elegans* males, we took an approach that focuses on social interaction. In a quadrant- format assay, ∼10 sexually naïve male “searchers” were allowed to interact with two classes of conspecific “targets” located in adjacent quadrants (Figure 1B). To prevent targets from migrating during the assay, they carried mutations in the muscle myosin *unc-54*.^39^ After placing searchers on the plate, we scored their locations at 30, 60, and 90 minutes to calculate a Mate Choice Index (MCI) (Figure 1B; see Materials & Methods). By focusing on the decision to approach and stay with potential mates, this assay captures the social interaction aspect of mate-choice behavior without considering progeny production, which is subject to the separate and potentially confounding influences of copulation efficiency and hermaphrodite fertility.

A fundamental feature assessed during mate-choice behavior is the sex of conspecifics. When we presented male searchers with adult targets of both sexes, they exhibited a strong preference for hermaphrodites, with roughly 75% of searchers localizing to the hermaphrodite-containing quadrants (mean MCI = 0.49 ± 0.23; Figure 1C). Males also showed a robust preference for hermaphrodites when the targets were L4 larvae (Figure 1D), even though L4s lack differentiated genitalia and cannot mate. However, males exhibited no preference with sexually immature mid-larval (L3) targets (Figure 1D), demonstrating that the production of these cues is developmentally regulated. We also directly compared the attractiveness of L4 to adult hermaphrodites. Males clearly preferred adults (Figure 1E), indicating that the cues produced by adult hermaphrodites are more attractive than those made by L4s.

Mate-choice behaviors typically differ by sex: most animals tend to choose mates of the opposite sex. Interestingly, we found that young adult hermaphrodites exhibited no apparent preference for interaction with males over hermaphrodites (Figure 1F). Though this experiment is unlikely to have the power to detect a small effect, this result is consistent with the idea that self-fertile hermaphrodites have more incentive to self-fertilize than to mate with males.^19^ Together, these findings demonstrate that males can robustly distinguish the sex and stage of potential mates using mate-choice cues presented in their native context and at physiologically relevant concentrations. Further, they show that the preference for interacting with hermaphrodites is a male-specific feature.

### Ascaroside pheromones, shaped by the sexual state of the intestine, contribute to mate choice decisions

To identify signals important for the sex-discrimination aspect of mate choice, we first considered the ascaroside pheromones. Several members of this family, particularly ascr#3 (also called “C9”) and ascr#8, are produced primarily by adult hermaphrodites and, as purified compounds, potently attract or retain males.^26, 40^ A related pheromone, ascr#10, is produced primarily by males and, in its purified form, attracts hermaphrodites.^41^ However, the contribution of endogenously produced ascarosides in male mate-choice decisions has not been carefully examined. To address this, we used targets carrying mutations in *daf-22*, an enzyme necessary for the synthesis of short-chain ascarosides including ascr#3, ascr#8, and ascr#10.^40, 42, 43^ Interestingly, males readily recognized the sex of *daf-22* adults, strongly preferring to interact with *daf-22* hermaphrodites over *daf-22* males (Figure 2A). However, this was not the case for L4 targets: males exhibited no sex preference when choosing between L4-stage *daf-22* males and hermaphrodites (Figure 2A). The simplest explanation of these findings is that ascaroside sex pheromones are produced by both L4 and adult hermaphrodites, but additional non-ascaroside signals are presented only by adults.

**Figure 2.**
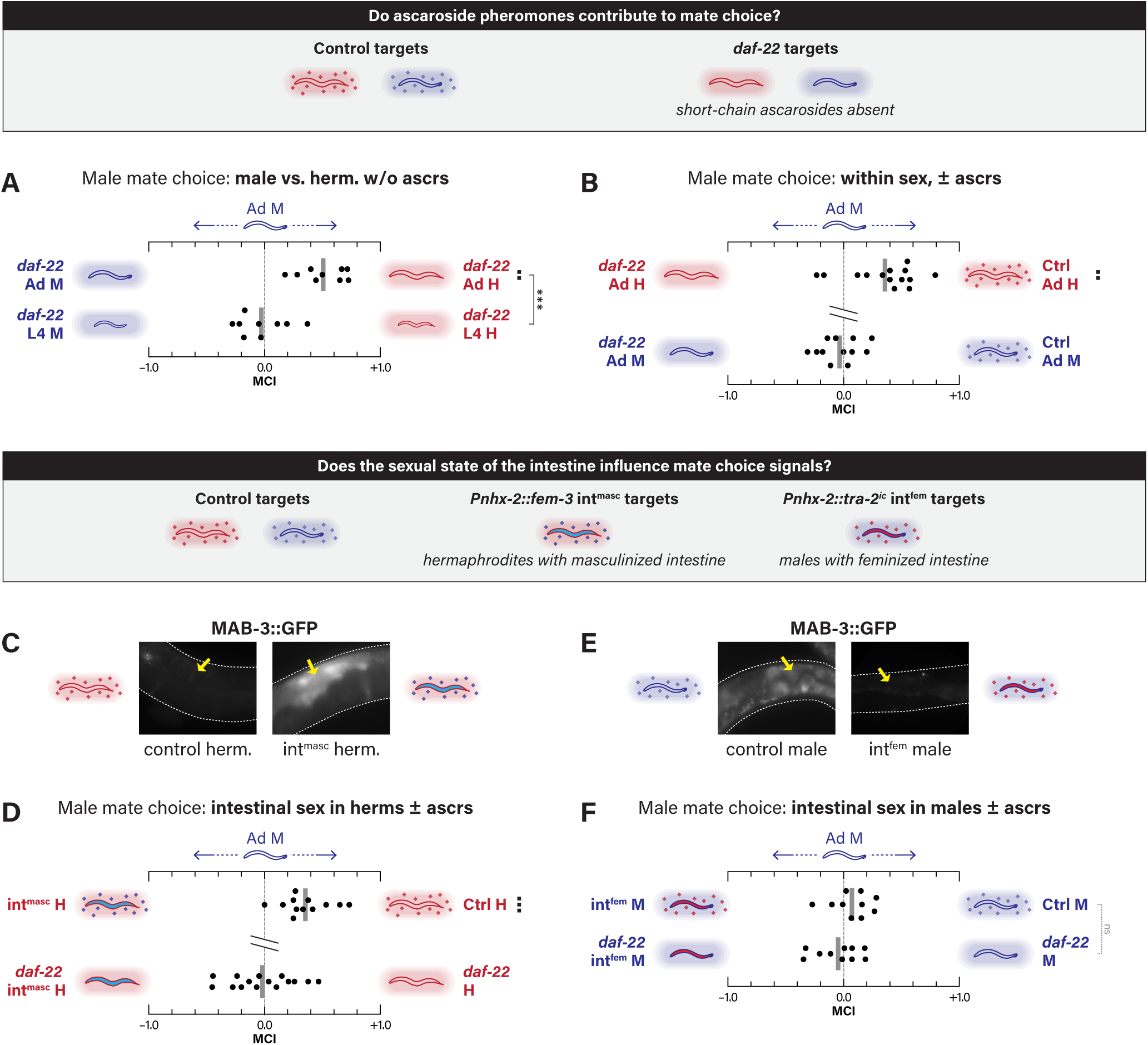
Ascaroside pheromones, whose sexual specificity is determined by the intestine, have central roles in sex and stage discrimination. (A, B) MCAs with control male searchers and *daf-22* targets that lack short-chain ascaroside production. (A) Between-sex MCAs with *daf-22* adults and L4 larvae. (B) Within-sex MCAs with *daf-22* vs. control adult hermaphrodites and males. The break in the vertical axis here and in subsequent panels below indicates that these experiments were not done in parallel, so we chose not to carry out a direct statistical comparison between them. (C, E) Epifluorescence images of *mab-3::GFP* in control and int^masc^ hermaphrodites (C) and in control and int^fem^ males (E). Dashed white lines outline the body; yellow arrows indicate the intestine. (D, F) Within-sex MCAs with control male searchers and control vs. sex- reversed intestine targets with or without short-chain ascarosides. (D) int^masc^ vs. control hermaphrodites in a wild-type and *daf-22* mutant background. (F) int^fem^ vs. control males in a wild-type and *daf-22* mutant background.

Although ascarosides are not necessary for males to determine the sex of adults, they do contribute to mate choice: males preferred to interact with control over *daf-22* adult hermaphrodites (Figure 2B), consistent with previous results using a slightly different assay.^35^ In contrast, when given control vs. *daf-22* male targets, males showed no apparent preference (Figure 2B). Thus, ascr#10 and other male-enriched ascarosides do not have strong male-aversive properties in this context.

ascr#3 and other short-chain ascarosides are produced by peroxisomal beta- oxidation in the *C. elegans* intestine.^44–47^ Sexual dimorphism in ascaroside production is thought to emerge from the regulated expression of peroxisomal enzymes in this tissue.^41^ To ask whether the sexual state of the intestine influences the attractiveness of a potential mate, we used “int^masc^” hermaphrodites, in which the sexual state of the intestine is masculinized through expression of the male sexual regulator *fem-3*.^48^ We found that int^masc^ hermaphrodites express the normally male-specific intestinal marker *MAB-3::GFP*,^49^ validating this manipulation (Figure 2C). Further, recent studies have shown that these animals produce male-typical ascaroside profiles.^50^ Consistent with this, we found that males robustly preferred control over int^masc^ hermaphrodite targets in the mate-choice assay (Figure 2D). This preference was completely dependent on *daf- 22* (Figure 2D), indicating that it arises from differential ascaroside production. Because *daf-22* is also expressed in the nervous system,^51^ we also tested neuro^masc^ targets (hermaphrodites carrying a previously validated pan-neuronal masculinization transgene^52^). However, with or without *daf-22* function, this manipulation had no apparent effect (Figure S1A). Thus, to mate-searching males, ascarosides report the sexual state of a potential mate’s intestine, but not its nervous system.

Genetic feminization of the male intestine would be expected to similarly sex- reverse patterns of ascaroside production, although this has not been tested directly. To create int^fem^ males, we expressed an activated form of the hermaphrodite sexual regulator *tra-2*^53^ in the intestine. As expected, *MAB-3::GFP* was repressed in the intestine of int^fem^ males (Figure 2E). However, males had no preference for int^fem^ over control males, regardless of *daf-22* genotype (Figure 2F). Thus, feminization of the male intestine may be insufficient to trigger production of hermaphrodite-typical pheromone profiles, or the effects of hermaphrodite pheromones produced by int^fem^ males may be overridden by unknown aversive cues produced by males. Together, these results indicate that the female state of the intestine is necessary to produce attractive ascarosides in hermaphrodites, but that it is not sufficient to make males attractive mates.

### The ability to copulate does not drive mate-choice decisions

Next, we considered roles for non-ascaroside cues in mate choice. Because a role for these is most apparent in adult targets (Figure 2A), we asked whether the hermaphrodite vulva, an adult-specific structure,^54^ is important in mate-choice behavior. However, males showed no detectable preference for controls over vulvaless *lin-39*^55^ adult hermaphrodites (Figure 3A). Therefore, neither the vulva itself, nor the ability to successfully copulate with the target, is a principal determinant of mate-choice behavior. However, when the influence of ascarosides was removed by comparing *daf- 22* against *daf-22; lin-39* targets, males displayed a weak preference for hermaphrodites with normal vulvae (Figure 3B). Thus, the vulva has a minor role whose influence becomes apparent only when stronger cues are removed. This could indicate that the vulva has an instructive or permissive role in generating a sensory signal detected by males; alternatively, an apparent preference might emerge from an increase in the time males spend with hermaphrodites during copulation itself, which is possible only with hermaphrodites with normal vulvae.

**Figure 3.**
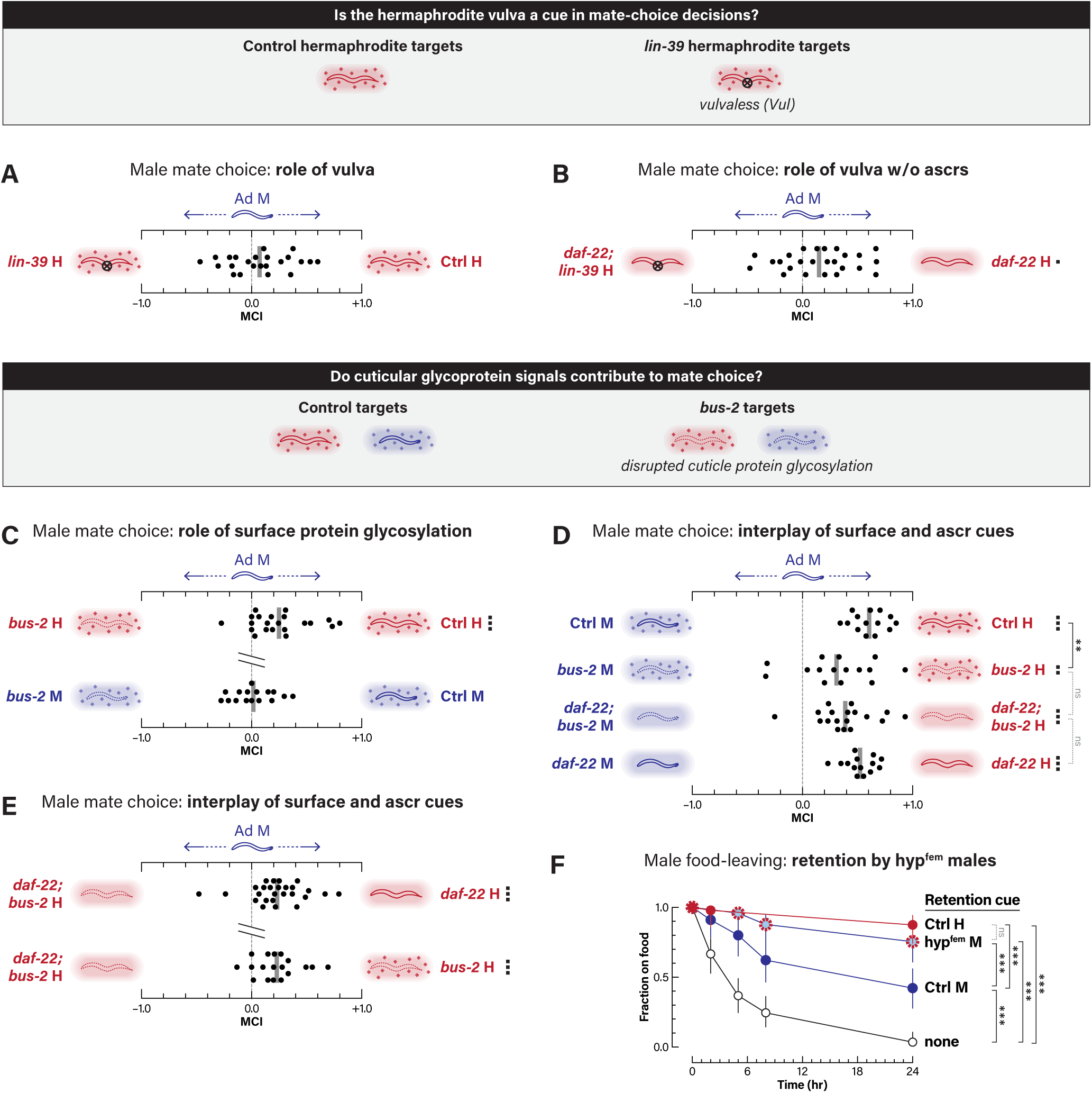
Morphological and surface cues also contribute to sex discrimination. **(A, B)** MCAs with control male searchers and (A) Vul *lin-39* vs. control hermaphrodites and (B) Vul *lin-39* vs. control hermaphrodites in the absence of short-chain ascarosides. **(C-E)** MCAs with control male searchers and (C) within-sex comparisons of *bus-2* vs. control hermaphrodites and males; (D) between-sex comparisons using control targets, *bus-2* targets, *daf-22; bus-2* targets, and *daf-22* targets; and (E) within-sex comparisons of *daf- 22; bus-2* vs. *daf-22* and *daf-22; bus-2* vs. *bus-2* targets. **(F)** Male food-leaving assay, measuring retention by different classes of targets: none, control males, control hermaphrodites, and hyp^fem^ males. Data points show the fraction of males remaining on a food spot harboring the indicated retention cue over 24 hr.

### The hermaphrodite cuticle provides mate-choice cues

Contact-associated cues, both chemical and mechanical, could play important roles in male mate-choice decisions.^38^ Previous work has established the importance of surface cues in species recognition during nematode mating.^56^ These signals also have roles in the initiation of male mating behavior,^32, 38^ in hermaphrodites’ ability to suppress male food-leaving behavior,^36, 37^ and in the detection of hermaphrodites by males in the context of sexual conditioning.^57^ Though the nature of these cues is not well understood, surface glycoproteins are likely to be involved, as males spend less time in contact with hermaphrodites carrying mutations in machinery responsible for the glycosylation of cuticle proteins.^32^

To ask whether surface glycoproteins have a role in mate recognition, we used animals carrying mutations in *bus-2*, a putative glycosyltransferase implicated in the O-linked glycosylation of surface proteins and the production of cues used by males to identify hermaphrodites.^32, 58^ Importantly, *bus-2* loss alters but does not abolish surface protein glycosylation,^58^ so these animals should not be thought of as entirely lacking these cues. Consistent with previous results,^32^ we found that males had a slight preference for interaction with wild-type over *bus-2* hermaphrodites (Figure 3C). Target *bus-2* activity had no apparent role in male-male interactions (Figure 3C), suggesting that male-specific patterns of surface glycosylation might not deter male-male interactions. *bus-2* likely functions in the hypodermis to promote hermaphrodite attraction, since hypodermal-specific expression rescued the *bus-2* attractiveness phenotype (Figure S1B), consistent with the reported expression of *bus-2* in hypodermal seam cells and the idea that it regulates the glycosylation of cuticle proteins.^58^

We next asked whether *bus-2-*depedent cues were important for sex discrimination. Interestingly, this ability was significantly reduced, though still intact, in a *bus-2* mutant background (Figure 3D). To probe the relative contributions of surface and ascaroside cues, we assayed mate choice with *daf-22; bus-2* hermaphrodite and male targets. MCI was not further reduced compared to either single mutant (Figure 3D), demonstrating that yet other cues are sufficient for sex discrimination in this context. We also compared mate choice with *daf-22* vs. *daf-22; bus-2* hermaphrodites and *bus-2* vs. *daf-22; bus-2* hermaphrodites. In both cases, the single mutants were significantly more attractive than the doubles (Figure 3E), indicating that each gene promotes attractiveness independently of the other, consistent with their distinct biochemical functions and sites of action. Thus, ascaroside and *bus-2-*dependent surface cues converge to optimize male mate choice.

The sex-specificity of surface cues might be instructed by the sexual state of the hypodermis, which synthesizes the worm’s cuticle. Genetic masculinization of the hermaphrodite hypodermis causes vulval rupture and death (our unpublished observations), making hyp^masc^ animals unsuitable targets in the mate-choice assay. Therefore, we asked instead whether feminization of the male hypodermis might make these animals more attractive to males. For these studies, we used an assay particularly sensitive to contact signals, the ability of hermaphrodites to retain males on a small food patch.^36, 37^ In the absence of conspecifics, males will readily abandon a food spot over a 24 hr period; however, the presence of hermaphrodites almost completely blocks this behavior (Figure 3F). As expected,^36^ we found that control males retained males to some degree; remarkably, however, hyp^fem^ males retained other males nearly as effectively as wild-type hermaphrodites (Figure 3F). Thus, the genetic sex of the hypodermis is a key determinant of the production of surface signals important for male mate-choice decisions.

### Male mate-choice decisions integrate volatile sex pheromones that signal self-sperm depletion

Beyond ascarosides, the vulva, and surface cues, another class of potential mate choice signals in *C. elegans* are the volatile sex pheromones (VSPs).^27, 29^ Some VSPs are abundantly produced by aged adult hermaphrodites after self-sperm depletion, thought to be an adaptation that reflects an increased motivation to attract mates.^27, 30^ Germline-defective mutant hermaphrodites mimic this effect, producing abundant VSPs even as young adults.^27^ While the molecular nature of VSPs is unknown, their production is likely independent of *daf-22* and they are therefore unlikely to be chemically related to ascarosides.^30^ Recent work has shown that males detect at least some VSPs using the putative chemoreceptor *srd-1*, which is expressed male-specifically in the sex-shared AWA olfactory neurons.^29^

To ask whether VSPs contribute to sex discrimination in young adults, we used *daf-22*; *bus-2* targets to minimize cue redundancy. We found that *srd-1* mutant males could distinguish the sex of *daf-22; bus-2* adults as well as control males (Figure 4A). Thus, *srd-1*-detected VSPs do not have a major role in this context, consistent with observations that young adult hermaphrodites produce negligible amounts of VSP.^29, 34^ The residual preference for hermaphrodites in these experiments could come from a variety of sources, including *bus-2*-independent surface cues, the hermaphrodite vulva, variant VSPs whose detection is independent of *srd-1*, long-chain ascarosides that are still produced in *daf-22* hermaphrodites, or entirely new classes of cues; this will require further study.

**Figure 4.**
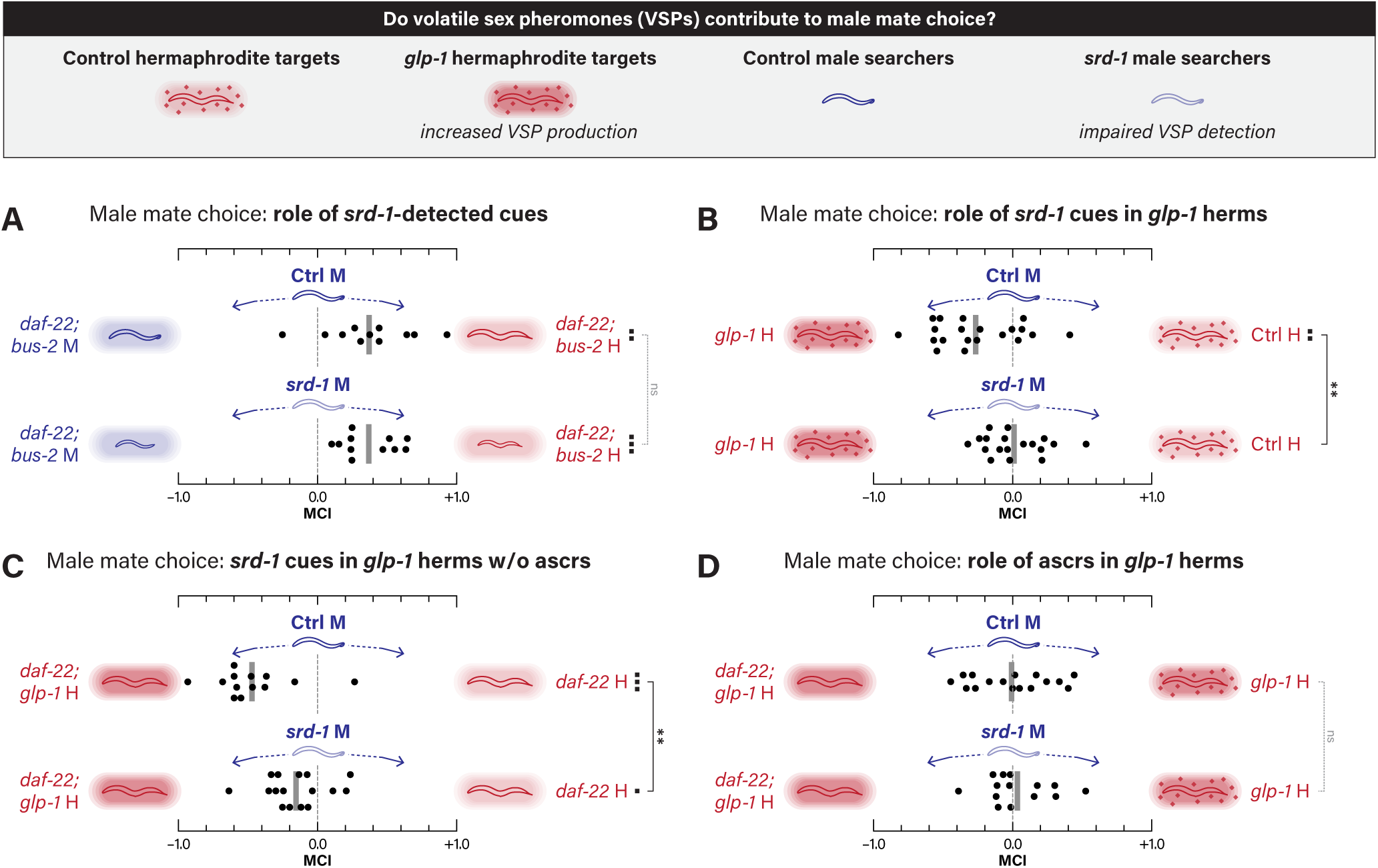
Volatile sex pheromones (VSPs) that signal fertility status are powerful mate choice signals. **(A-D)** MCAs with wild-type and *srd-1* mutant male searchers. (A) Between-sex MCAs with *daf-22; bus-2* targets. (B) Within-sex MCAs with *glp-1* vs. control targets. (C) Within-sex MCAs with *daf-22; glp-1* vs. *daf-22* targets. (D) Within-sex MCAs with *daf-22; glp-1* vs. *glp-1* targets.

We were unable to directly examine older hermaphrodites in this assay because the *unc-54* mutation used to immobilize targets causes animals to retain eggs and become unhealthy before they are depleted of self-sperm. To circumvent this, we used *glp-1* mutants, in which germline loss causes young hermaphrodites to increase VSP production.^27^ We found that males showed a significant preference for interacting with *glp-1* over wild-type hermaphrodites (Figure 4B), suggesting that VSPs can have a strong influence on mate choice. This effect was absent at 15°C, the permissive temperature for *glp-1*,^59^ confirming its specificity (Figure S1C). The increased attractiveness of *glp-1* hermaphrodites was completely dependent on *srd-1* function in males (Figure 4B), demonstrating that VSPs detected by *srd-1* are the sole mediator of this property. When we removed the contribution of ascarosides, VSPs appeared to play an even larger role and, surprisingly, their effect was only partially dependent on *srd-1* (Figure 4C). This suggests that young adult *glp-1* hermaphrodites produce additional cues, perhaps including *srd-1*-independent VSPs, and that the activity of some of these is apparent only when the stronger influence of ascarosides is removed.

To compare the properties of VSPs and ascarosides, we allowed males to choose between *daf-22; glp-1* and *glp-1* hermaphrodite targets. Here, both target classes produce high VSP but only one produces ascarosides. Surprisingly, *daf-22*-dependent cues had no apparent role in this context (Figure 4D), suggesting that the long-range potency of volatile VSP cues overwhelms the short-range attractive effects of ascarosides. Interestingly, this was also the case when males lacked *srd-1* (Figure 4D), further supporting the idea that *srd-1* detects only a subset of VSPs. An alternative possibility, that sterile *glp-1* hermaphrodites produce fewer attractive (or more repellent) ascarosides, is unlikely given recent findings that the gonad has no role in short-chain ascaroside synthesis.^50^ Together, our results indicate that mate-choice decisions involving wild-type young adult hermaphrodites do not depend on VSP signals, but that these molecules are likely to be powerful cues promoting preferential mating with older hermaphrodites. Further, our results confirm a key role for *srd-1* in detecting natively produced VSPs but also implicate additional chemoreceptor(s) in detecting these cues.

### Ascaroside signals allow males to assess the nutritional status of potential mates

Beyond sex and developmental stage, other features of potential mates may be important indicators of their reproductive fitness and could therefore be useful for males to assess. For *C. elegans* males, knowing a hermaphrodite’s nutritional status could be especially relevant, as nutrient-deprived hermaphrodites produce low-quality oocytes and have small broods.^60^ To determine whether males can recognize nutritional stress in a potential mate, we allowed them to choose between well-fed and fasted adult hermaphrodite targets. With 4-hr fasted targets, males showed no discernable preference (Figure 5A). However, males clearly preferred well-fed to 28-hr fasted targets (Figure 5A), demonstrating that males can detect more severe starvation in a potential mate. To ask whether this preference stems from differences in ascaroside biosynthesis, we carried out the same experiment using *daf-22* targets. Unexpectedly, males exhibited a robust preference for well-fed over 4-hr fasted *daf-22* hermaphrodites, and this effect was absent at 28 hr (Figure 5B). These data suggest two conclusions. First, the aversive effects of long-term fasting could be mediated by ascarosides. Consistent with this, L1 larvae starved for ∼12 hr have been shown to increase production of the aversive ascaroside osas#9 and decrease levels of the male attractant ascr#3 compared to well-fed L1s.^43, 61^ Second, the rapid production of aversive cues by 4-hr starved *daf-22* hermaphrodites might reflect a sensitization to nutrient deprivation arising from their altered fatty acid metabolism,^46, 62^ potentially promoting the production of non-ascaroside aversive compounds. Together, these results indicate that males assess the nutritional status of potential mates and that ascaroside and non- ascaroside chemical signals may both have roles in this process.

**Figure 5.**
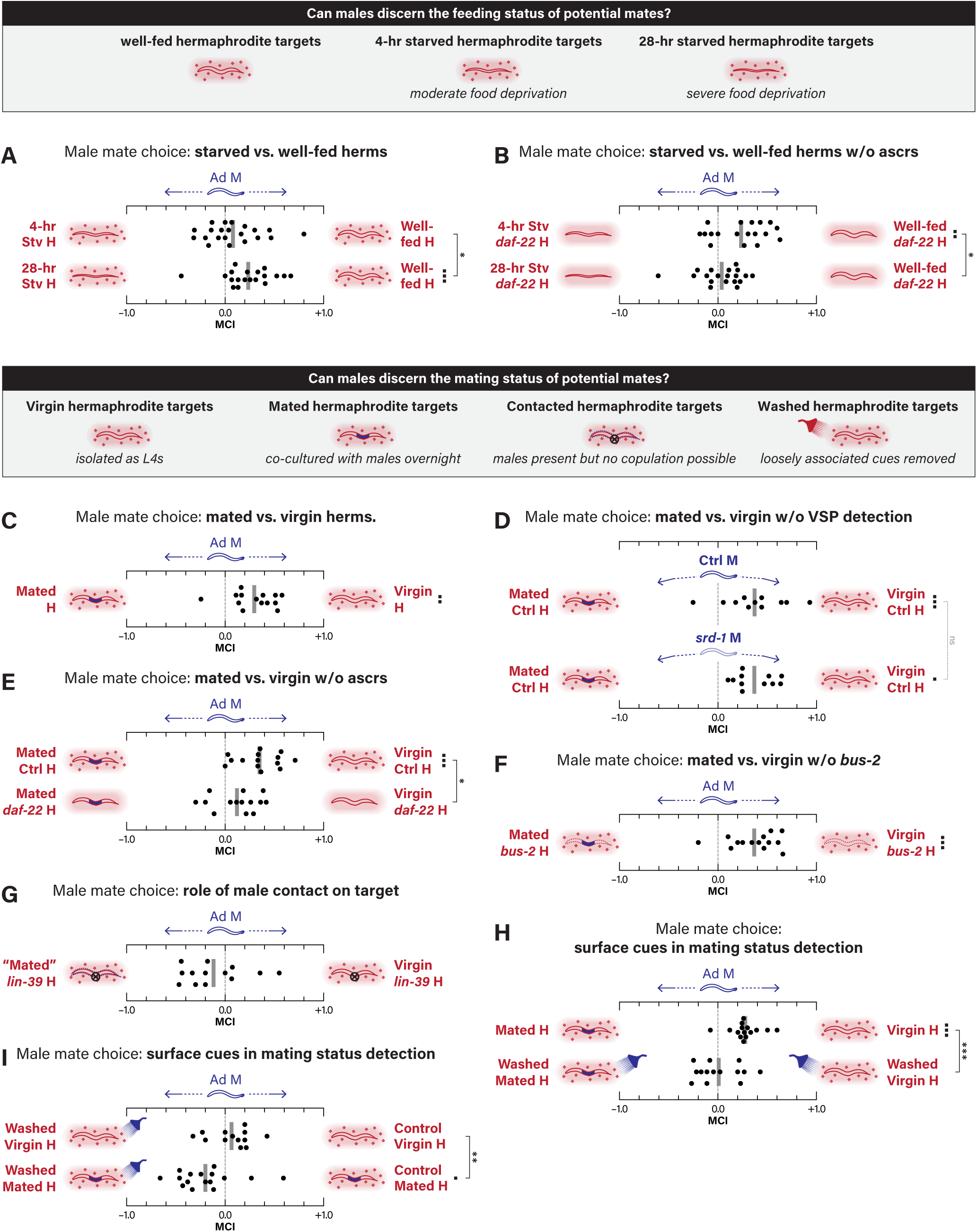
*C. elegans* males assess target nutritional state and reproductive history in mate-choice decisions. **(A,B)** MCAs with control male searchers and food-deprived hermaphrodite targets. (A) 4-hr and 28-hr nutrient-deprived vs. control hermaphrodite targets. (B) 4-hr and 28-hr nutrient-deprived vs. control hermaphrodite targets, with all targets carrying a *daf-22* mutation. **(C-I)** MCAs with male searchers and virgin or previously mated hermaphrodite targets. (C) Control male searchers with previously mated vs. virgin young adult hermaphrodites. (D) Control and *srd-1* mutant male searchers with mated vs. virgin hermaphrodites. (E) Control male searchers with mated vs. virgin hermaphrodites, either in a wild-type or *daf-22* background. (F) Control male searchers with mated vs. virgin *bus-2* hermaphrodites. (G) Control male searchers with Vul *lin-39* targets previously exposed to males (“mated”) vs. control targets exposed to *unc-54* males than cannot mate (“virgin”). (H) Control male searchers with mated vs. virgin targets that had not (control) or had been briefly washed before testing. (I) Control male searchers with washed vs. control targets, either virgin or mated.

### Multiple cues allow males to detect the mating history of potential mates

Mating history is another feature of hermaphrodite physiology with important reproductive fitness consequences. Even though both virgin and mated young adult hermaphrodites are gravid, males might prefer to avoid choosing a previously mated hermaphrodite if virgins are available, particularly since male sperm have a fitness advantage over hermaphrodite self-sperm.^63^ Further, males mating with virgin hermaphrodites might actively reduce their mates’ attractiveness to deter competitors. Consistent with this, males are slightly more attracted to conditioned media produced by virgin adult hermaphrodites compared to mated ones^64^ and mating decreases the strength of VSP cues produced by germline-defective *fog-2* hermaphrodites.^27^

To ask whether mating status is incorporated into mate-choice decisions, we let males choose between young adult hermaphrodites that had been co-cultured for 16 hours with young adult *him-5* males or instead with sterile *unc-54; him-5* males, allowing us to control for exposure to male pheromones, which can dramatically alter hermaphrodite physiology.^65, 66^ Males displayed a clear preference for virgin over mated hermaphrodites, indicating that they can robustly detect this quality and integrate it into their behavior (Figure 5C). Because previous work has suggested or indicated that fertility status can alter VSP production,^27, 30, 34^ we asked whether the VSP receptor *srd- 1*^29^ was necessary for this preference. *srd-1* mutant males still clearly preferred virgin hermaphrodites, and their preference index did not differ from those of control males (Figure 5D). However, because *srd-1* may not be responsible for all VSP detection, this does not rule out a role for VSPs in this behavior.

To consider the possibility that ascaroside or surface cues produced by mated hermaphrodites might differ from those of virgins, we asked whether males could distinguish mating status in targets lacking *daf-22* or *bus-2*. Interestingly, we found that the virgin-vs.-mated MCI was significantly lower with *daf-22* than with control targets (Figure 5E), likely indicating that the production of ascaroside sex pheromones by hermaphrodites changes upon mating. In contrast, loss of *bus-2* function in targets had no apparent effect on males’ ability to detect their fertilization status (Figure 5F).

Because we observed a trend (*p* = 0.079) in males’ ability to detect mating status in the absence of ascarosides (Figure 5E), we tested the possibility that males might detect a cue deposited by a previous mate on the surface of the hermaphrodite cuticle. In many nematode species, including some natural *C. elegans* isolates, males deposit a mating plug on their mates, which could decrease the attractiveness to other males.^67^ Although males of the laboratory strain N2 have lost this ability,^67^ they could still leave evidence of their encounter with a hermaphrodite while their tail is scanning the hermaphrodite’s body. To test this, we used Vul *lin-39* hermaphrodites that had been previously co-cultured with males. Males still attempt copulation with Vul hermaphrodites, repeatedly scanning the length of their body with their tails. However, we did not detect a statistically significant preference when males chose between *lin-39* hermaphrodites that had or had not previously been co-cultured with fertile males (Figure 5G). Thus, vulva-scanning behavior alone does not appear to leave a cue that decreases a hermaphrodite’s attractiveness.

Males could also deposit cues on the hermaphrodite body during copulation itself;^68, 69^ further, copulation has been associated with damage to the hermaphrodite cuticle near the vulva, which could cause the release of a signal.^70^ To evaluate these possibilities, we washed hermaphrodites briefly before placing them on the assay plate.

Under these conditions, preference toward virgin hermaphrodites was abolished (Figure 5H), strongly suggesting that a surface cue accounts for this behavior. To explore this further, we directly compared washed vs. control animals. Washing virgin hermaphrodites had no effect on their attractiveness to males (Figure 5I). However, with previously mated hermaphrodites, males clearly preferred washed over control targets (Figure 5I). From this, we infer that an aversive surface-associated cue decreases the attractiveness of mated hermaphrodites. Extracellular vesicles deposited in and on hermaphrodites by males during mating^68, 71^ are interesting candidates for mediating this function (see Discussion).

### The male state of sex-shared neurons is essential for responses to multiple classes of sex discrimination cues

We next investigated the mechanisms by which the male nervous system detects and interprets signals guiding mate-choice decisions. Because the preference for interacting with hermaphrodites is a male-specific feature, we focused on the neuronal mechanisms that could underlie this dimorphism. The CEM neurons are the only class of male-specific head sensory neurons in *C. elegans*;^72^ previous work has shown that they can detect synthetic ascarosides.^26, 73^ We found that *ceh-30* mutant males, which lack CEM neurons,^74, 75^ could still efficiently distinguish target sex (Figure S2A), consistent with the finding that ascaroside cues are not essential for sex discrimination (Figure 2A). We assessed the role of CEMs in ascaroside detection more directly by comparing males’ preference for wild-type over *daf-22* adult hermaphrodites. On average, *ceh-30* males still behaved comparably to control males; however, there was a marked increase in the variability of their behavior (F-test, *p* = 0.0016) (Figure S2B). This suggests that ascaroside detection by CEMs is important for robust and consistent mate-choice decisions, but that CEM function is not necessary for the response to physiologically relevant, natively produced ascarosides. *ceh-30* males robustly discerned the sex of *daf- 22* mutants, indicating that the CEMs are not essential for ascaroside-independent sex discrimination (Figure S2C). We also tested males lacking the TRPP-class polycystin channel *pkd-2*, important for the function of the CEMs as well as the RnB and HOB sensory neurons of the male tail.^76^ Though severely impaired in their ability to respond to physical contact with hermaphrodites,^76^ *pkd-2* males were fully able to distinguish target sex in the absence of ascarosides (Figure S2D).

These results led us to ask whether male-specific features of sex-shared neurons and circuits are important for mate-choice decisions. Consistent with this possibility, males carrying a pan-neural feminization transgene (*Prab-3::tra-2^ic^*, “neuro^fem^” males)^53^ completely lost their preference for hermaphrodites (Figure 6A). Moreover, masculinizing the hermaphrodite nervous system was sufficient to masculinize behavior: *Prab-3::fem-3* neuro^masc^ hermaphrodites^28, 77^ showed a clear preference for interaction with hermaphrodites over males (Figure 6A). Because *Prab-3* is not active early enough in development to alter precursor lineages, neuro^masc^ hermaphrodites do not possess male-specific neurons; instead, sex-shared neurons adopt male-typical physiological and anatomical features.^28, 77^ Thus, male-specific tuning of shared circuits is sufficient to establish a preference for interaction with hermaphrodites.

**Figure 6.**
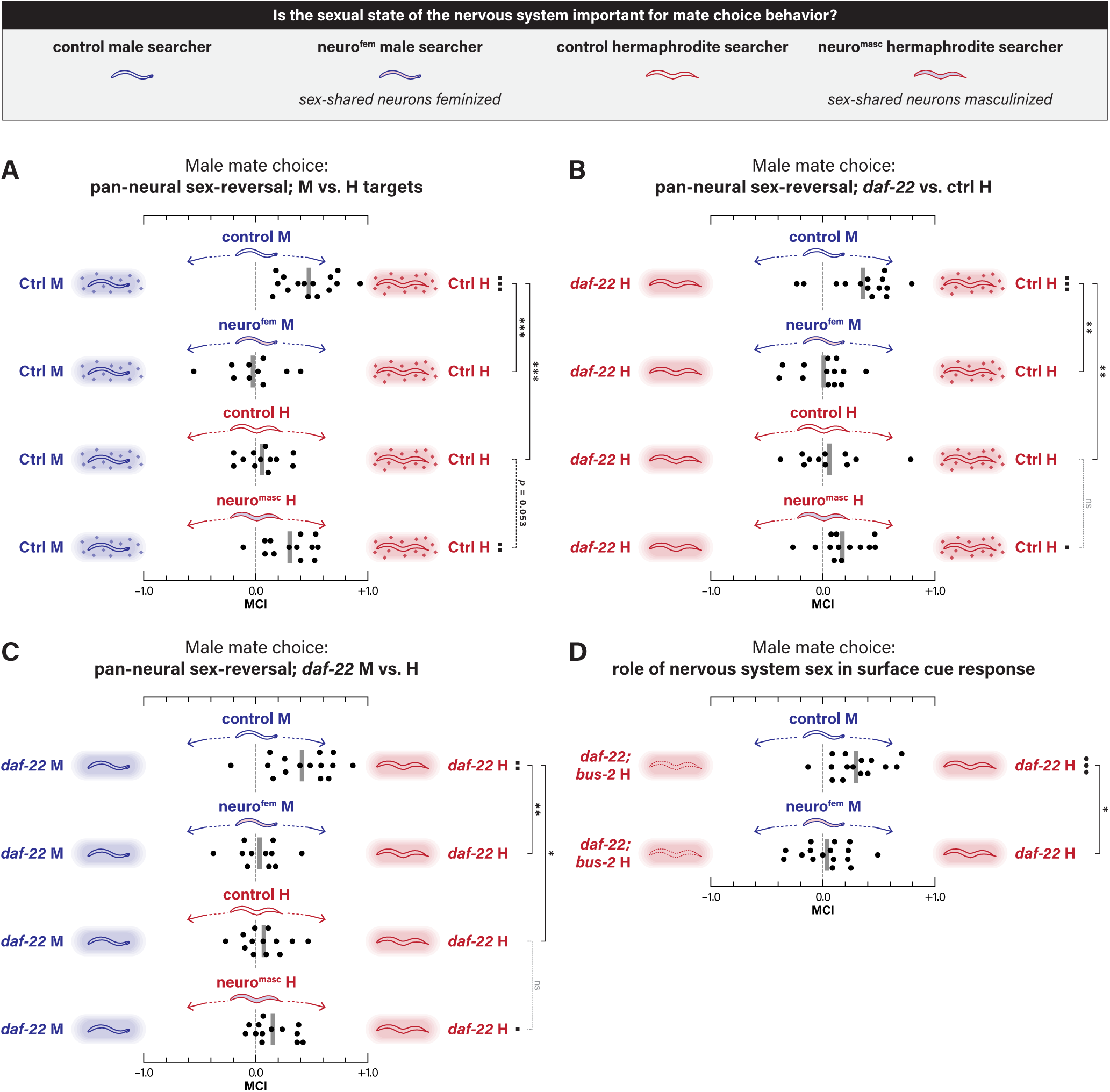
The male state of shared neural circuits is necessary and sufficient for aspects of male mate-choice behavior. **(A-C)** MCAs with control male, neuro^fem^ male, control hermaphrodite, and neuro^masc^ hermaphrodite searchers. (A) Between-sex MCAs with control male and hermaphrodite targets. (B) Within-sex MCAs with *daf-22* and control hermaphrodite targets. (C) Between-sex MCAs with *daf-22* male and hermaphrodite targets. **(D)** MCA with control and neuro^fem^ male searchers and ***daf-22; bus-2* vs. *daf-22*** hermaphrodite targets.

We next asked how the sexual state of the nervous system influences searchers’ responses to different classes of mate-choice cues, focusing first on ascarosides.

Neuro^fem^ males lost their preference for control over *daf-22* hermaphrodites (Figure 6B), confirming a requirement for the male state of shared circuits in ascaroside detection.^35^ The behavior of neuro^masc^ hermaphrodites was not different from controls, though these animals did display a slight and statistically significant preference for interacting with control over *daf-22* hermaphrodites (Figure 6B). Previous work has shown that neuro^masc^ hermaphrodites are robustly attracted to purified ascarosides;^35^ the fact that these animals do not exhibit stronger mate preference behavior suggests that other male-specific features may be important for the robust responses to natively produced hermaphrodite ascarosides.

We also explored the mechanisms underlying males’ response to non-ascaroside cues. Previous findings have indicated that the detection of hermaphrodite surface cues is mediated by male-specific sensory structures.^38^ Surprisingly, the ability to distinguish the sex of *daf-22* targets was abolished in neuro^fem^ males (Figure 6C), revealing a requirement for sex-shared circuits in the response to these cues. Consistent with this model, the behavior of neuro^masc^ hermaphrodites, which do not possess male-specific sensory structures, was statistically indistinguishable from that of control hermaphrodites (Figure 6C). Interestingly, however, these animals did display a slight preference for interacting with hermaphrodite targets (Figure 6C). This indicates that the male state of the nervous system is necessary for the detection and/or interpretation of non-ascaroside cues, and, in hermaphrodites, is sufficient to elicit a small response to these.

To more specifically ask whether sexual modulation of shared circuits is important for the response to surface cues, we tested males’ ability to distinguish between *daf-22; bus-2* and *daf-22* targets. Pan-neuronal feminization completely abolished the preference for *daf-22* targets (Figure 6D). Thus, male-specific features of shared neurons—either shared sensory neurons or interneurons downstream of male- specific sensory neurons—are important for detecting or interpreting information on the surface of potential mates.

### The sex-shared ADF and ASH sensory neurons have key roles in males’ response to natively produced ascaroside sex pheromones

Sex-specific sensory tuning is a prominent feature of the *C. elegans* nervous system,^28, 29, 35, 52, 77–79^ prompting us to explore roles for specific sensory neurons in detecting natively produced mate-choice cues. Feminization of all ciliated sensory neurons (sens^fem^) significantly disrupted males’ ability to distinguish *daf-22* from control targets (Figure 7A). To assess the roles of individual sensory neurons, we assayed this behavior in a series of strains carrying single-neuron genetic ablations. The sensory neurons ASK and ADL can detect ascaroside pheromones,^26, 78^ but ASK ablation revealed no apparent defect in males’ ability to read ascaroside cues (Figure 7B). ADL-ablated displayed a marginally reduced MCI (*p* = 0.093) and higher variance in behavior (F-test, *p* = 0.039), so we cannot draw a strong conclusion regarding a role for this neuron in the response to native ascarosides (Figure 7C). We also tested AWB-, ASI-, and ASE/AWC- ablated males, revealing no apparent defects in this behavior (Figure 7D).

**Figure 7.**
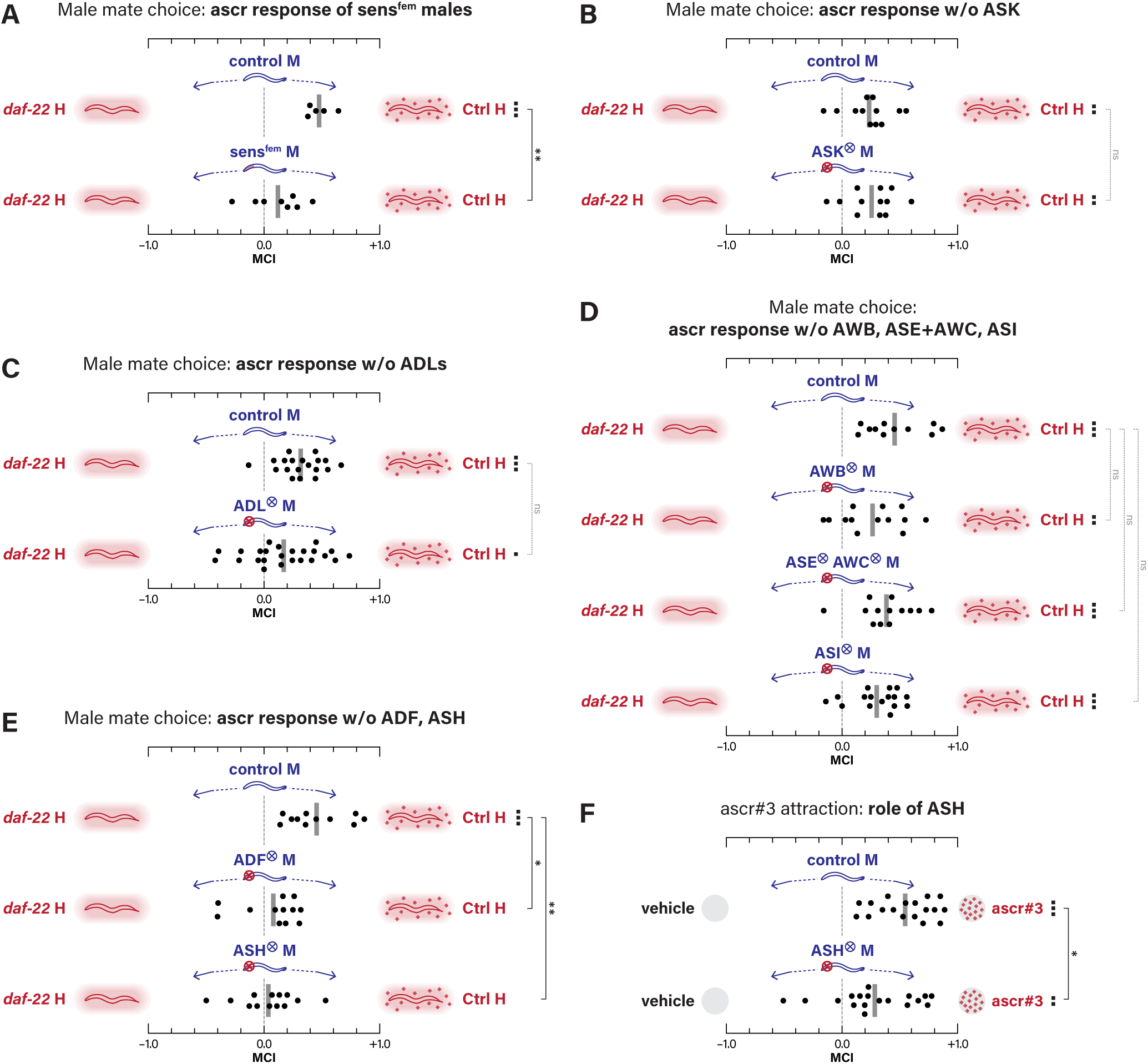
The sex-shared ADF and ASH neurons have key roles in ascaroside-mediated mate-choice decisions. **(A-E)** MCAs with *daf-22* vs. control hermaphrodite targets. (A) MCAs with control and sens^fem^ male searchers. (B) MCAs with control and ASK-ablated male searchers. Strains used in this panel contained *him-8(e1489)* rather than *him- 5(e1490)*. (C) MCAs with control and ADL-ablated male searchers. (D) MCAs with control searchers and those with ablation of AWB; AWE and AWC; or ASI. (E) MCAs with control, ADF-ablated, and ASH-ablated male searchers. **(F)** Quadrant chemotaxis assay for ascr#3 attraction using control or ASH-ablated males.

Consistent with the role of the shared ADF sensory neuron in generating attraction to purified ascr#3,^35^ ADF ablation severely impaired the preference for control over *daf-22* hermaphrodites (Figure 7E). ADF was not essential for sex discrimination, either with control or *daf-22* targets (Figure S3A, B), consistent with the redundancy of signal used in this behavior. Unexpectedly, ablation of the polymodal nociceptive neuron ASH also disrupted males’ preference for control over *daf-22* targets (Figure 7E, S3C). Because ASH has not been implicated in ascaroside-evoked behavior, we tested ASH-ablated males in a quadrant-format ascaroside attraction assay. Indeed, these animals exhibited a marked decrease in ascr#3 attraction (Figure 7F), demonstrating that ASH is important, though not essential, for the response to this pheromone. ASH was not required for males to distinguish target sex in control or *daf- 22* animals (Figure S3D, E), again supporting the idea that multiple, redundant signals contribute to sex discrimination. Interestingly, recent work has shown that sex differences in synaptic connectivity reduce ASH’s contribution to avoidance behavior and permit robust mating in males.^80^ Our results raise the intriguing possibility that reconfiguration of ASH physiology and/or connectivity allow it to mediate a response to appetitive cues in males.

## DISCUSSION

While copulatory behavior in *C. elegans* males has received significant attention,^38, 81^ far less is known about how males make decisions about when to mate. Even the simplest aspect of this decision – assessing the sex and stage of potential mates – has not been systematically examined in the context of natively presented cues. Here, using a competitive social interaction assay that reflects behavior evoked by endogenously produced mate-choice cues, we identify a sex-specific behavioral program in which males use diverse sensory cues to assess multiple aspects of the reproductive fitness of potential mates. Because *C. elegans* males tend to initiate contact-response behavior with any object they encounter, a key determinant of mate-choice behavior must be the navigational decisions that bring males into proximity with optimal mates.

Further, since males use a combination of volatile, soluble, and contact-dependent cues, the physical nature of these signals determines a behavioral sequence guiding mate- choice behavior (Figure 8A). Together with previous findings, our results lead to an integrated framework for these decisions that incorporates diverse, multimodal stimuli, the nature of the information provided by these stimuli, and the neuronal mechanisms that underlie male-specific responses to them.

**Figure 8.**
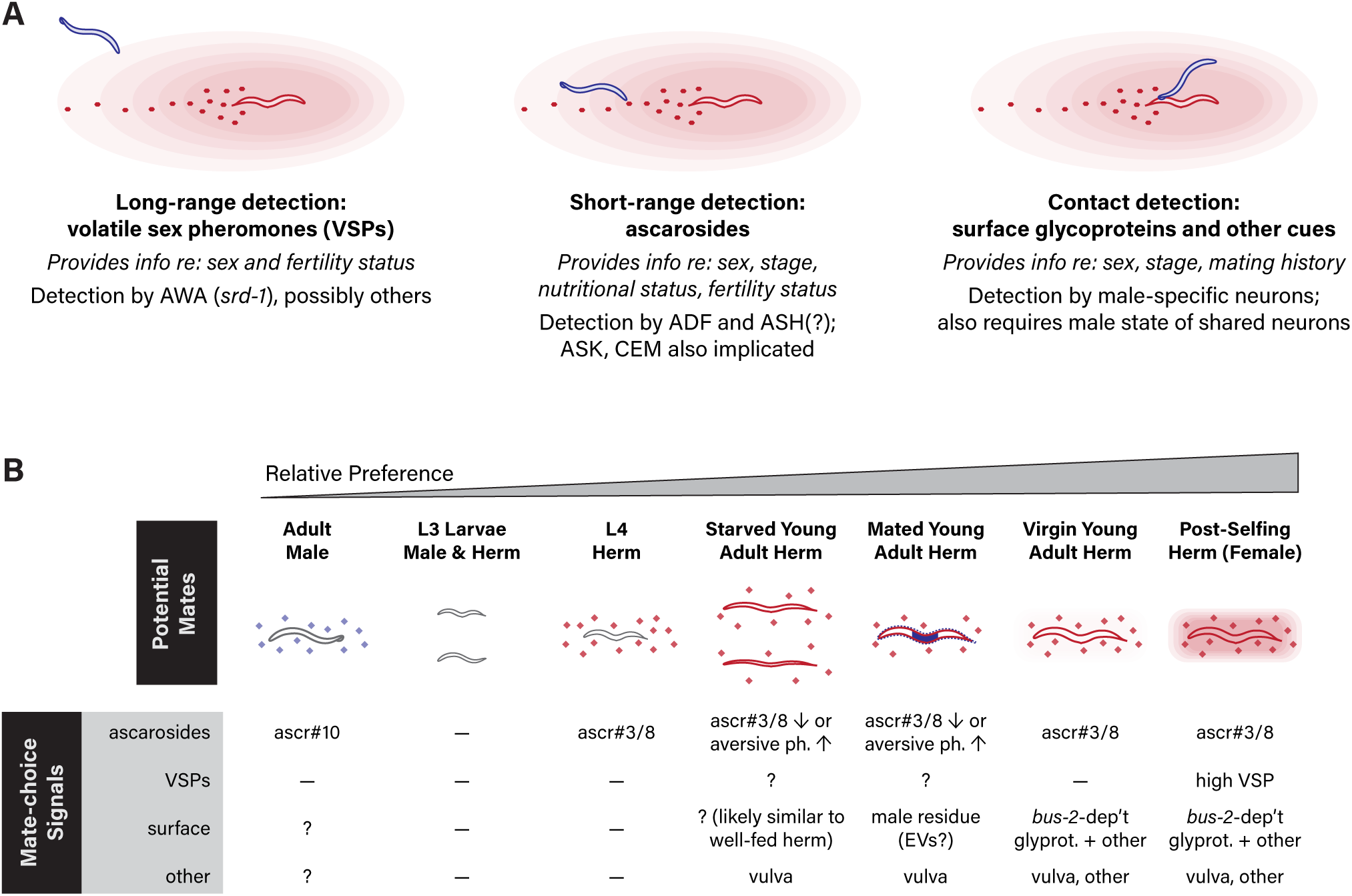
*C. elegans* males to assess multiple features of potential mates using a hierarchical behavioral strategy that incorporates multimodal sensory cues. (A) The hierarchy of mate-choice behavior is dictated by the nature of these diverse signals and the mechanisms by which they are detected. In this scenario, a hermaphrodite has been migrating to the right and has recently transitioned from roaming to dwelling behavior. A trail of ascaroside-containing deposits is indicated. As volatile cues,^27, 29^ VSPs likely attract males from a distance; this is probably most relevant for older, sperm-depleted hermaphrodites “advertising” for males. Less-diffusible ascaroside cues are thought to be deposited as frass and are likely non-uniform in distribution.^83^ These may allow males to identify regions in which hermaphrodites have recently been present and could trigger a transition to a local-search-like state.^35^ Detection and evaluation of surface cues likely happens during the initial phase of mating behavior, as male-specific neurons in the male tail contact the mate’s body during scanning behavior.^38^ Each of these three phases of mate-detection carries allows the male to assess specific subsets of the features of potential mates. (B) A table summarizing various categories of potential mates, ordered according to male preference. For each category, the role of different classes of mate-choice signals is indicated. “—” indicates that this signal is absent or does not play a detectable role in mate-choice. “?” indicates that the status of this signal is unknown.

In many settings, the first salient mate-choice cue detected by males is likely to be long-range olfactory signals from the VSP class of sex pheromones (Figure 8). Our results, together with others,^27, 34^ indicate that well-fed, young adult hermaphrodites produce negligible amounts of these cues. However, we find that VSPs are essential for male preference for young *glp-1* hermaphrodites, which mimic the increased attractiveness of older, sperm-depleted hermaphrodites.^27, 30, 34^ VSPs are detected in part by the sex-shared AWA olfactory neurons via the receptor *srd-1*;^29^ however, our studies also suggest that *C. elegans* might produce additional VSPs whose detection is independent of *srd-1*. VSPs are primarily thought to signal sperm depletion, though it is possible that they could also be used by younger hermaphrodites in contexts that increase their motivation for outcrossing, *e.g.*, pathogen exposure.^82^ Identifying the molecular nature of these pheromones and the mechanisms that regulate their production will be important future goals for the field.

At closer range, ascaroside pheromones, particularly ascr#3 and ascr#8, likely dominate mate-choice decisions. These cues are not volatile; rather, evidence suggests that they are deposited in discrete packets during defecation and are only weakly diffusible.^83^ As such, they could signal the presence of a nearby hermaphrodite to a male, potentially by triggering a local search-like state of motor behavior. Ascaroside biosynthesis is modulated not only by sex but also by developmental stage and feeding status;^21–25^ consistent with this, our data indicate that males use ascarosides to distinguish not only sex but also adults from young (L3) larvae. The sexual information provided by ascarosides seems to be specified exclusively by the sexual state of the intestine, indicating that the master sexual regulator *tra-1*, possibly through its target *mab-3*,^49^ modulates metabolic pathways in this tissue. Further, our results indicate that males’ ability to distinguish feeding state and mating history in a potential mate is also dependent at least in part on ascarosides. Multiple neurons, both sex-specific and shared, have been implicated in ascaroside detection. In the assay used here, the male- specific CEMs have an apparently relatively minor contribution. Instead, the sex-shared ADF and ASH neurons are essential for the response to natively presented hermaphrodite ascarosides. A key role for male-specific features of ADF in ascaroside detection has previously been demonstrated;^35^ however, to our knowledge, a role for ASH in sex pheromone attraction has not been described. Whether ASH has sex-specific properties that enable it to detect or process pheromone signals will be an important question for future studies. Interestingly, recent work has shown that the nociceptive function of ASH is blunted in males due in part to sex differences in synaptic connectivity.^80^ Our results suggest that ASH might be repurposed in males, allowing it to contribute to appetitive responses.

The final phase of male mate-choice behavior depends on contact-dependent cues, but the nature of these cues and the mechanisms by which they are detected remains poorly understood. Interestingly, recent work has found that the mechanical cue of body stiffness is an important determinant of the recognition of hermaphrodites by males (C.-H. Chen et al., pers. comm.). We find that the glycosylation of surface proteins plays an important role in males’ ability to discriminate the sex of potential mates, consistent with previous work showing that males are less responsive to *bus-2* mutant hermaphrodites.^32^ Our results demonstrate that their sex-specificity of surface cues is determined by the genetic sex of the hypodermis itself, but the chemical nature of these dimorphisms and the ways in which they are controlled by the sexual regulator *tra-1* remains unclear. Previous work has established that these cues are detected by male-specific sensory neurons in the tail;^38, 84^ our results also indicate a requirement for the male state of shared circuitry in males’ ability to respond to them.

Other kinds of surface-associated cues are also important in male mate-choice. Our data suggest that detection of the hermaphrodite vulva has a secondary role in promoting sustained male-hermaphrodite interaction, providing a means to distinguish not only sex but also developmental stage. Interestingly, surface-associated cues also appear have a role in allowing a male to assess the mating history of adult hermaphrodites. Extracellular vesicles (EVs) are intriguing candidates for mediating this effect, as males are known to shed neuronally derived EVs in the vicinity of the vulva during mating behavior.^71^ However, exposure to male-derived EVs has been shown to enhance, rather than suppress, male mating behavior.^68^ Further work will be necessary to dissect the role of male-deposited EVs and other cues on male mate-choice decisions.

Our studies show that multiple steps of mate-choice behavior depend critically on the male state of sex-shared neurons. Interestingly, neur^fem^ males remain fertile and sire cross-progeny (data not shown), suggesting that the male state of shared neurons is not essential for copulatory behavior. Instead, male-specific features of sex-shared neurons may facilitate other male-typical behaviors, including the ability to assess mate quality, that involve reconfiguring sex-shared circuits. One of the most intriguing findings from the recent description of the connectome of the adult *C. elegans* male has been the extensive sexual dimorphism in synaptic connectivity among shared neurons,^85^ which arises through sex-specific pruning and synaptogenesis that rewires multiple sensory circuits.^80, 86^ Our results indicate that successful integration of inputs from male-specific neurons requires the male state of shared circuitry, and that male-specific synaptic and neuromodulatory configurations of sex-shared neurons are important for implementing male-specific sensory integration and behavioral states.

A rich view of the reproductive behavior of the *C. elegans* male, particularly in its native habitats, remains enigmatic. *C. elegans* reproduces primarily by self-fertilization and the species shows some hallmarks of “selfing syndrome,” a degradation of traits that promote sexual reproduction.^18, 87^ However, population genetic studies reveal evidence of sporadic outcrossing in wild populations.^18^ Thus, male-mate choice behavior could be an important determinant of the contribution of cross-fertilization to fitness, particularly under times of stress. Our results demonstrate that males prefer to interact with virgin over previously mated hermaphrodites, and with well-fed over nutrient-deprived hermaphrodites. Males can likely assess other aspects of hermaphrodite state—recent studies have shown that hermaphrodites exposed to the pathogen *Pseudomonas* have an increased mating frequency,^82^ but that males avoid mating with hermaphrodites infected with Orsay virus.^88^ Each of the steps of mate-choice behavior we have described here are also likely plastic; understanding how they are influenced by a male’s physiological status and reproductive experience, as well as the behavioral and physiological features of potential mates, will be exciting areas for future work.

## MATERIALS AND METHODS

### Experimental Model

All *C. elegans* strains were cultured using *E. coli* OP50 and NGM agar as described.^89, 90^ All strains were grown and assayed at 20°C, except for those containing *glp-1* mutations; for these, *glp-1* parents were cultured at 15°C from egg to young adult stage to permit germline development and then transferred to 25°C to generate germline-less *glp-1* targets, which were assayed at 20°C. All relevant control strains experienced identical temperature shifts. All target strains contained the mutation *unc- 54(e190)* to prevent animals from migrating on the assay plate. All strains were derived from N2 and, unless otherwise noted, contained *him-5(e1490)* to increase the frequency of spontaneous males in self-fertilizing populations.

### Molecular Biology and Construction of Transgenic Strains

Sex-reversal transgenes were generated using the Multisite Gateway Cloning System (Invitrogen). The intestine-specific *Pnhx-2* promoter was used to drive expression of *fem-3(+)* or *tra-2(ic)* to masculinize or feminize this tissue, respectively. *Pdpy-7* was used to feminize the hypodermis. Transgenic animals were generated using injection mixes containing 100 ng/μL of the co-injection marker *Psulp-3::gfp* and 20-50 ng/μL of the *fem-3(+)* or *tra-2(ic)* expression construct.

### Mate Preference Assay

One day prior to the assay, L4 animals (both searchers and targets) were picked to sex-segregated plates. To prepare assay plates, quadrant boundaries were marked on the bottom of unseeded 6-cm NGM plates with a custom-made stamp. The quadrant area was a square (3 cm on each edge) divided into four zones with a 1-cm diameter circle in the center. *E. coli* OP50 fresh culture (50 µl) was spread within the 3 cm square and plates were incubated at 20°C overnight. The next day, four target animals were placed in the center of each quadrant (Fig. 1B) at *t* = -30 mins. All targets carried *unc- 54(e190)* to limit their mobility. At time *t* = 0, ten searcher animals were placed in the center circle; their positions were then recorded at 30 min intervals (t = 30 min, 60 min, and 90 min). Plates on which target animals moved outside their quadrant boundaries (an uncommon occurrence) were censored. We calculated Mate Preference Index values for each time point [MCI = (# in “A” quadrants – # in “B” quadrants)/ (# in “A” quadrants + # in “B” quadrants)] and averaged these to obtain a final MCI. Thus, in the graphs shown, each data point represents the behavior of 10 searchers over 90 minutes.

### Preparation of fasted targets

To prepare fasted targets, *unc-54* or *unc-54; daf-22* hermaphrodites were fed until the young adult stage and then hand-picked into a drop of sterile water freshly loaded on the surface of bacteria-free NGM agar. After soaking briefly (2-3 min), they were hand-picked into bacteria-free NGM agar plate to initiate the fasting process (*t* = 0). At *t* = 4 hr, half of these animals were moved into assay plates to be used as 4hr- fasted targets. The remainder continued to fast on the bacteria-free agar until *t* = 28 hr, when they were moved into assay plates to be used as 28h-fasted targets. No bacterial growth was observed on the fasting plates at this time. To create well-fed controls, similar steps were performed at the same time points, except worms were placed on OP50-seeded NGM agar plates instead of bacteria-free ones after rinsing.

### Preparation of mated and virgin targets

To prepare mated targets, 50 young adult *him-5* males and 50 L4 *unc-54* or *unc- 54; daf-22* hermaphrodites were hand-picked to an NGM agar plate containing a small (∼1 cm diameter) OP50 bacteria lawn. These plates were incubated for 24 hr at 20°C, after which time males will have mated with the vast majority of hermaphrodites. To prepare control virgin targets, we incubated L4 hermaphrodites with *unc-54; him-5* males, which cannot mate^14^ but still expose hermaphrodites to male pheromones.

### Quadrant Assay with synthetic ascr#3

This assay was performed as described previously,^35^ using 1 µM purified ascr#3, a generous gift of F. Schroeder (Boyce Thompson Institute and Cornell University).

### Mate-retention using the food-leaving assay

The ability of a potential mate to suppress food-leaving behavior was assayed as previously described.^36, 37^

### Transgenic animals

Worm total genomic DNA was purified from mixed-stage DR466 with a commercial kit (Qiagen catalog number 51304) as the template of *dpy-7* promoter amplification. Worm total RNA was purified from mixed-stage DR466 with a commercial kit (Qiagen catalog number 74134), followed by *in vitro* cDNA synthesis (Invitrogen catalog number 18080-051), as the template of *bus-2* (isoform a) cDNA amplification.

Other reagents used for the construction of expression plasmids include the MultiSite Gateway Three-fragment Vector Construction Kit (Invitrogen Catalog number 12537- 023), DH5α competent cell (ThermoScientific catalog number 18265017). Plasmid sequences were confirmed by Sanger sequencing (Genewiz Inc, NJ, USA). Standard microinjection procedures were used to generate transgenic animals with the desired plasmid and a fluorescent co-injection marker at a total DNA concentration of 100 ng/μL.

### Microscopy

Animals were prepared for epifluorescence microscopy using standard procedures. DIC and fluorescence images were acquired using a Zeiss Axioplan II. Images were obtained using consistent exposure times and were not subjected to brightness or contrast manipulations.

### Statistical analysis

For each tester-target condition, we carried out a Wilcoxon signed-rank test to ask whether the MCI differed from the null hypothesis that males had no preference for one class of targets over the other (*e.g.,* MCI = 0). The associated *p* values are indicated with black squares to the right of each next to each row: ▪, *p* ≤ 0.05; ▪▪, *p* ≤ 0.005; ▪▪▪, *p* ≤ 0.001. To compare MCI values between different classes of searchers, we used a Mann-Whitney test (to compare two genotypes) or a Kruskal-Wallis test with Dunn’s correction (to compare more than two genotypes). Asterisks indicate *p* values associated with these tests: \**p* ≤ 0.05; \*\**p* ≤ 0.005; \*\*\**p* ≤ 0.001. For clarity, the brackets in each graph indicate all comparisons made, including those whose outcomes were not statistically significant (labeled “n.s.”). In cases where the variance in individual MCI values appeared to be exceptionally high, we carried out F-tests to determine whether MCI variance differed significantly from paired controls. Except for the food- leaving assay, all statistical tests were carried out using Prism 9.5 (GraphPad Software, LLC). For the food-leaving assay, PL (probability of leaving per worm per hr) values for each genotype were calculated using R (www.rproject.org) to fit the censored data with an exponenCal parametric survival model, using maximum likelihood. The hazard values obtained were reported as the PL values. To estimate PL values, s.e.m. and the 95% confidence intervals, worms from each experimental treatment were pooled across replicas and contrasted against controls using maximum likelihood.

### Key Resources Table

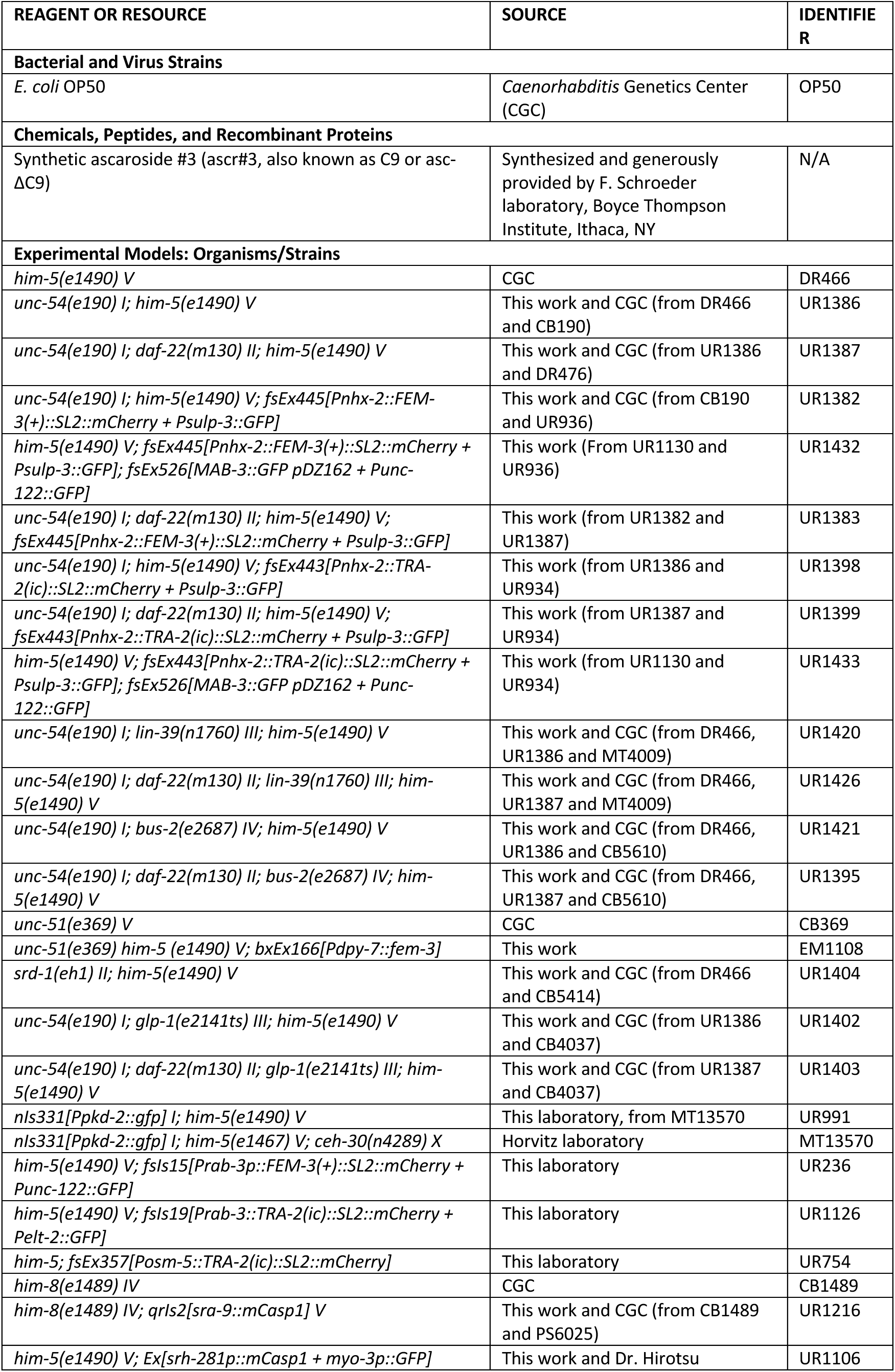
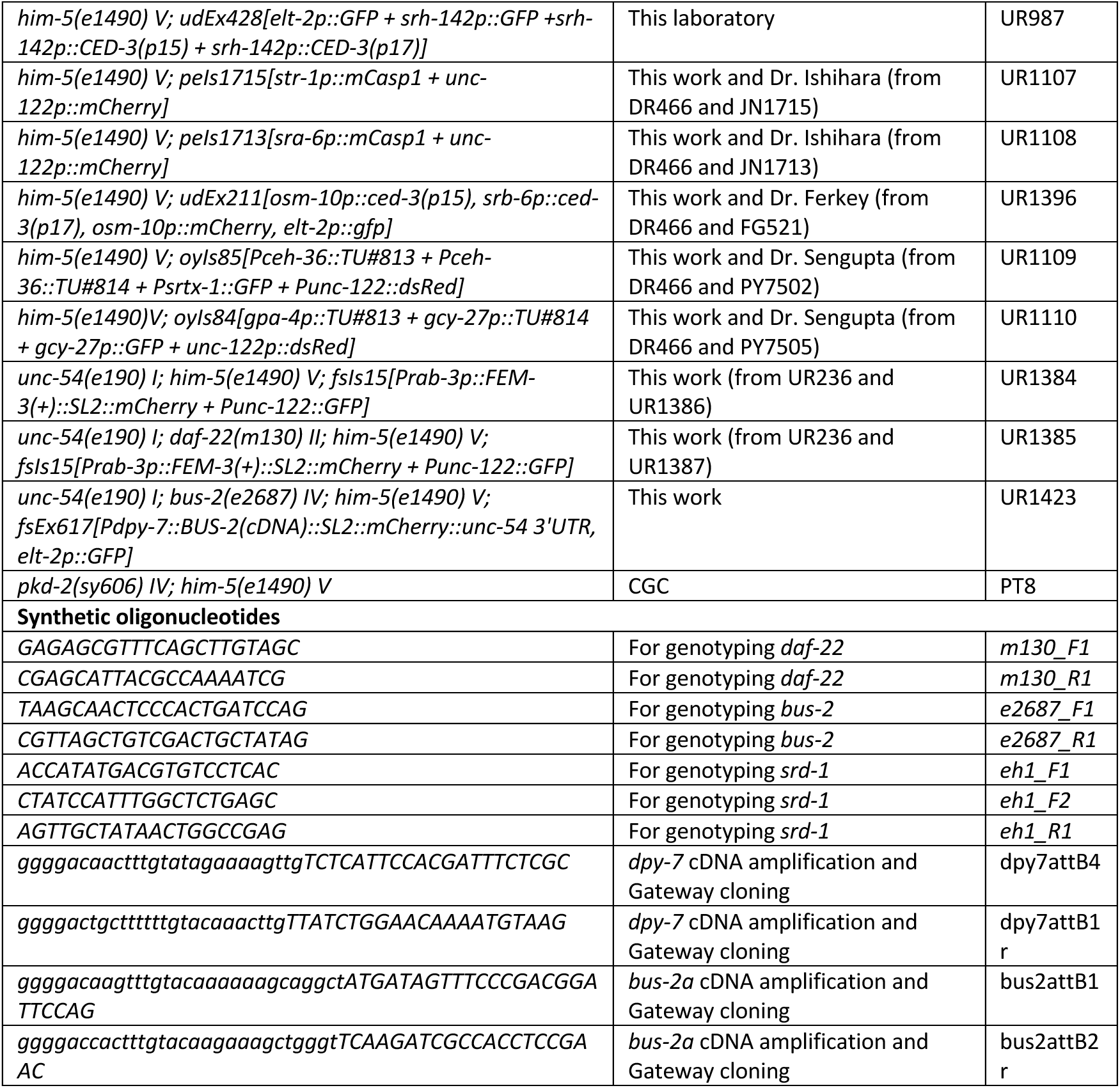

## ACKNOWLEDGEMENTS

We thank current and past members of the Portman lab, the University of Rochester Invertebrate Biology Group, the Western New York Worm Group, Scott Emmons, Maureen Barr, and Gabriella Sterne for discussion and critical feedback. J.L. thanks Dr. Yong Yu at Xiamen University for providing a supportive working environment. We are grateful to D. Ferkey, T. Hirotsu, T. Ishihara, and P. Sengupta for sharing strains and to F. Schroeder for purified ascarosides. Some strains used in this work were provided by the *Caenorhabditis* Genetics Center, which is funded by NIH Office of Research Infrastructure Programs (P40 OD010440). We thank WormBase, an essential resource for these studies, which is supported by Grant U41 HG002223 from the National Human Genome Research Institute at the NIH, the UK Medical Research Council, and the UK Biotechnology and Biological Sciences Research Council. These studies were funded by NIH R01 GM130136, R01 GM140415, and R35 GM148349 to D.P. and by Leverhulme Trust project grant RPG-2018–287 to A.B.

## AUTHOR CONTRIBUTIONS

Conceptualization: J.L., D.P., and A.B.; Investigation: J.L., A.B. and R.M.; Writing — Original Draft: J.L. and D.P.; Writing — Review & Editing: D.P.; Supervision: D.P.; Funding Acquisition: D.P. and A.B.

---

**Figure S1.**
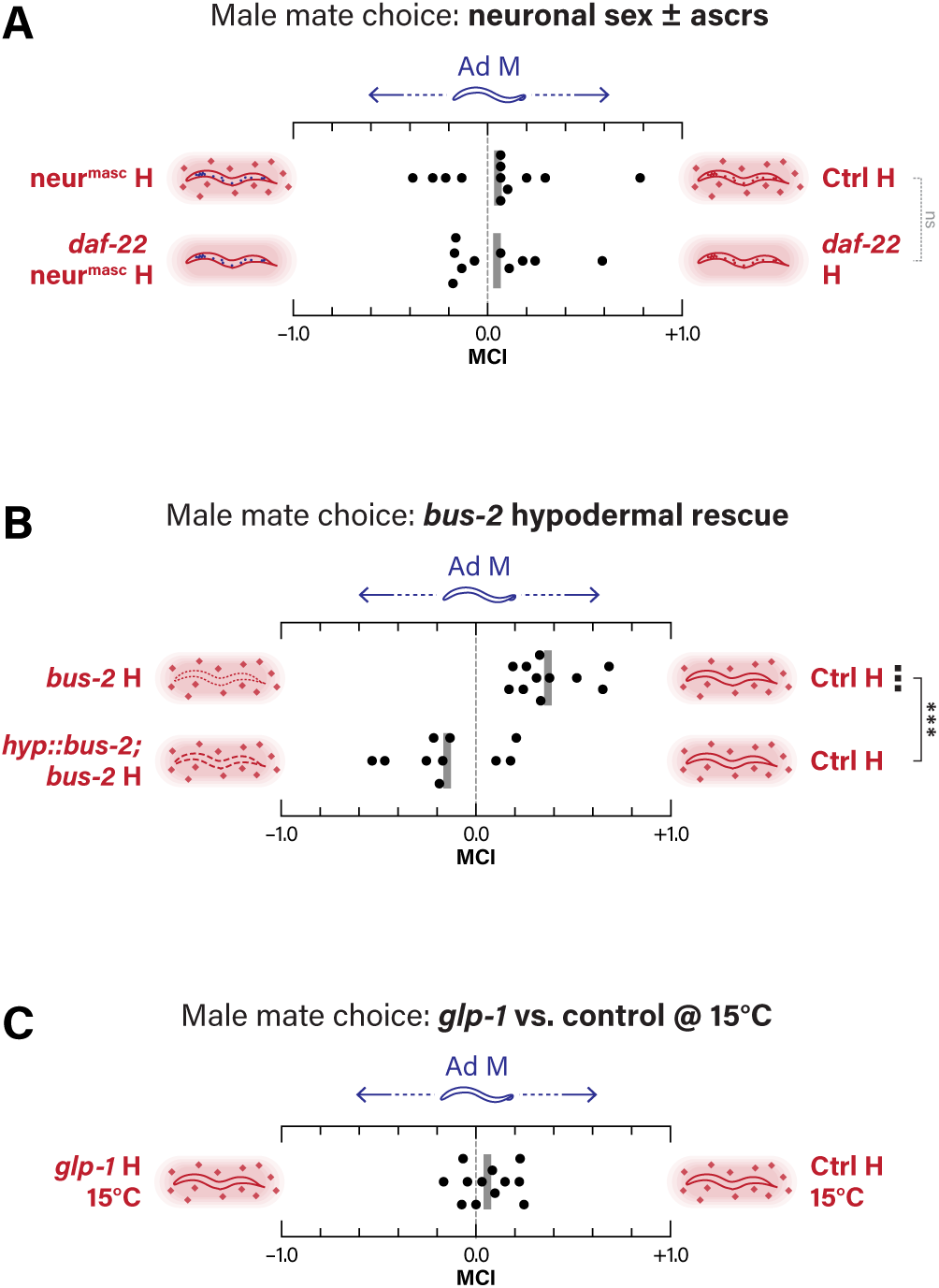
Related to Figures 2, 3, and 4. **(A-C)** Within-sex MCAs with control male searchers. (A) neur^masc^ vs. control hermaphrodite targets, in wild-type or *daf-22* backgrounds. (B) *bus-2* vs. control targets and rescued (*hyp::bus-2; bus-2*) vs. control targets. (C) *glp-1* vs. control targets grown at the permissive temperature of 15°C.

**Figure S2.**
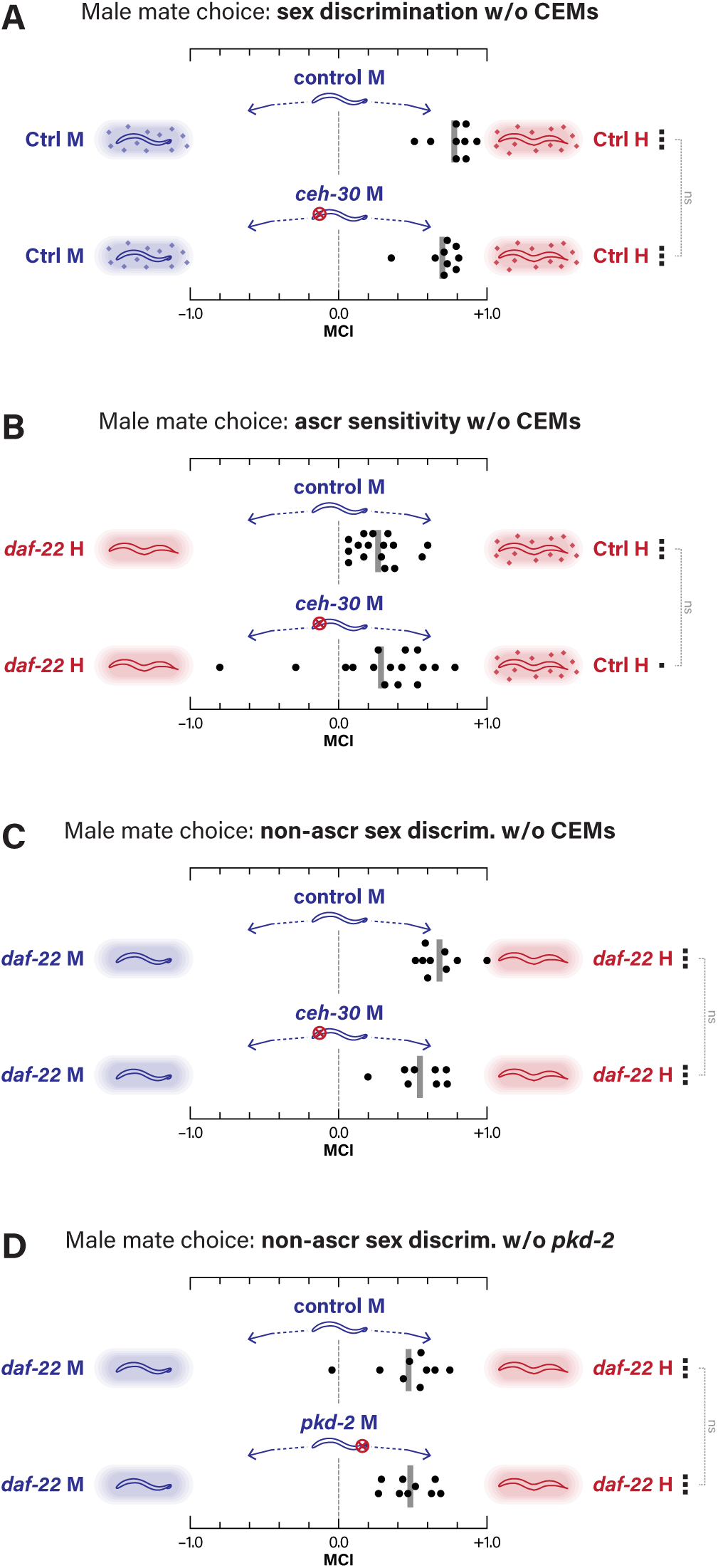
Related to Figure 6. **(A)** Between-sex MCAs with control and *ceh-30* male searchers with control targets. **(B)** Within-sex MCAs with control and *ceh-30* male searchers and *daf-22* vs. control targets. **(C)** Between-sex MCAs with control and *ceh-30* male searchers with *daf-22* targets. **(D)** Between-sex MCAs with control and *pkd-2* male searchers with *daf-22* targets.

**Figure S3.**
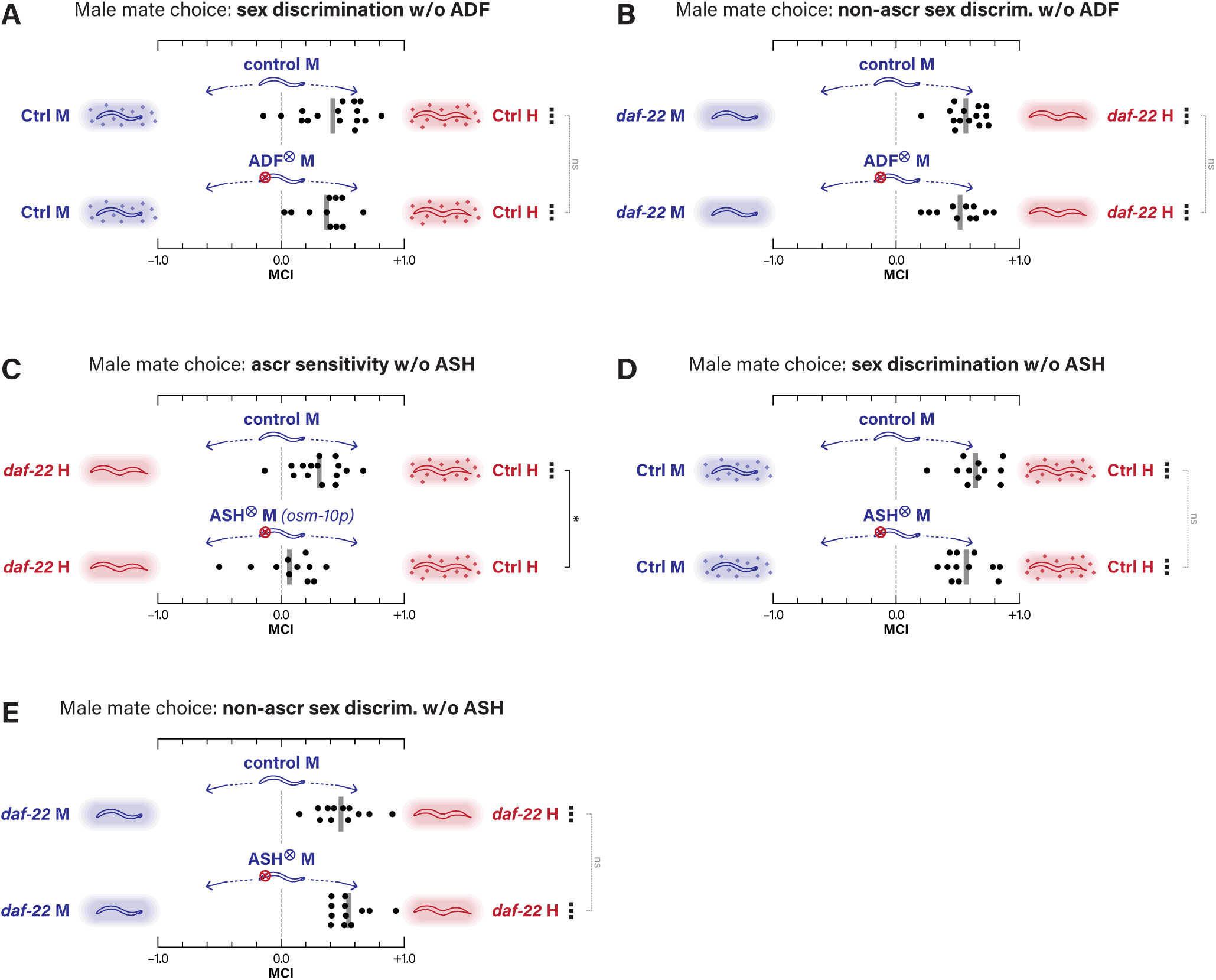
Related to Figure 7. **(A)** Between-sex MCAs with control and ADF-ablated male searchers with control targets. **(B)** Between-sex MCAs with control and ADF- ablated male searchers with *daf-22* targets. **(C)** Within-sex MCAs with control and ASH- ablated male searchers (using a *Posm-10* transgene) with *daf-22* vs. control targets. **(D)** Between-sex MCAs with control and ASH-ablated male searchers with control targets. **(E)** Between-sex MCAs with control and ASH-ablated male searchers with *daf-22* targets.

